# Deciphering Multiway Multiscale Brain Network Connectivity: Insights from Birth to 6 Months

**DOI:** 10.1101/2025.01.24.634772

**Authors:** Qiang Li, Zening Fu, Hasse Walum, Masoud Seraji, Prerana Bajracharya, Vince Calhoun, Sarah Shultz, Armin Iraji

## Abstract

Converging evidence suggests that understanding the human brain requires more than just examining pairwise functional brain interactions. The human brain is a complex, nonlinear system, and focusing solely on linear pairwise functional connectivity often overlooks important nonlinear and higher-order relationships. Infancy is a critical period marked by significant brain development that could contribute to future learning, health, and life success. Exploring higher-order functional relationships in the brain can provide insight into brain function and development. To the best of our knowledge, there is no existing research on multiway, multiscale brain network interactions in infants. In this study, we comprehensively investigate the interactions among brain intrinsic connectivity networks (ICNs), including both pairwise (pair-FNC) and triple relationships (tri-FNC). We focused on an infant dataset collected between birth and six months, a critical period for brain maturation. Our results revealed significant hierarchical, multiway, multiscale brain functional network interactions in the infant brain. These findings suggest that tri-FNC provide additional insights beyond what pairwise interactions reveal during early brain development. The tri-FNC predominantly involve the default mode, sensorimotor, visual, limbic, language, salience, and central executive domains. Notably, these triplet networks align with the classical triple network model of the human brain, which includes the default mode network, the salience network, and the central executive network. This suggests that the brain network system might already be initially established during the first six months of infancy. Interestingly, tri-FNC in the default mode and salience domains showed significantly stronger nonlinear interactions with age compared to pair-FNC. We also found that pair-FNC were less effective at detecting these networks. The present study suggests that exploring tri-FNC can offer additional insights beyond pair-FNC by capturing higher-order nonlinear interactions, potentially yielding more reliable biomarkers to characterize developmental trajectories.

## 1 Introduction

The human brain is organized in a hierarchical, multiscale manner that facilitates efficient information processing, which is essential for maintaining functional brain network interactions [1, 2, 3, 4]. These interactions reflect the core principles that allow brain networks to specialize for specific functions. These principles involve mechanisms that coordinate brain activity across different scales of organization [5]. During infancy, this developmental period is characterized by the discovery of the underlying rules that govern network interactions. By examining multiscale, multiway interactions during this phase, we can gain valuable insights into how these mechanisms drive the emergence of higher-order interactions and the specialization of brain networks. Notably, this intricate process of network specialization and integration is particularly pronounced during the early developmental stage [6]. Specifically, the first six months represent a critical period when the brain undergoes substantial growth and reorganization, laying the foundation for the complex network interactions observed in later stages of life [7, 8, 9]. A significant aspect of this developmental phase involves uncovering the rules that govern network interactions within functional connectomes [10]. Gaining insight into how these processes evolve during the perinatal and early postnatal periods is essential for understanding the mechanisms behind both typical and atypical brain developments [7, 8, 11, 12].

Previous research using resting-state functional magnetic resonance imaging (rsfMRI) has shown that changes in functional connectivity during early human development are mainly driven by the maturation of primary brain systems, with relatively less impact on higher-level regions such as the default mode, salience, frontoparietal, and executive control networks [13, 14, 15, 16]. One main reason for this is that by the end of the first year, primary sensorimotor and auditory networks have developed to closely resemble their adult counterparts, whereas higher-level networks, which undergo considerable development and continue to develop [17]. Studies suggest that whole-brain connectivity efficiency improves significantly by one year and reaches a stable level by age 2 [18, 15]. After this period, brain development is thought to be characterized mainly by the reorganization and remodeling of established major functional networks, including the refining and optimizing of neural connections and the shaping of functional brain architecture [19]. Furthermore, by two years of age, previous studies suggest that the default network begins to show some resemblance to that observed in adults, involving regions such as the medial prefrontal cortex, the posterior cingulate cortex/retrosplenial, the inferior parietal lobule, the lateral temporal cortex, and the hippocampus [16, 18]. Furthermore, to explore brain network interactions during early brain development, functional connectivity is widely used to estimate these interactions. Pairwise metrics, such as Pearson correlation and mutual information, are frequently applied to assess both linear and nonlinear pairwise functional connectivity [20, 21, 22]. However, brain network interactions extend beyond pairwise connections, involving multiway interactions that these traditional metrics are unable to capture [20, 23, 24, 25, 26, 27].

To address these limitations, a metric derived from information theory, called total correlation [28], can be used to capture high-order interactions. Studies have demonstrated that total correlation provides more information than pairwise metrics and can enhance the diagnosis of brain disorders [23, 29, 30, 31]. In addition, previous studies often extract blood-oxygen-level-dependent (BOLD) signals from brain regions using predefined brain atlases with specific constraints. This approach is limited because it relies on fixed atlases, which may overlook subject specificity and cannot guarantee that the predefined regions represent the same functional entities [32, 33]. A more suitable alternative would be a data-driven method that identifies subject-specific, functionally homogeneous units [33, 34, 35]. One such technique is Independent Component Analysis (ICA), which identifies intrinsic connectivity networks (ICNs) directly from the BOLD signal without relying on predefined brain atlases or specific anatomical regions [36, 37, 38]. This allows for the discovery of brain networks based on the data itself, rather than imposing predefined constraints.

Therefore, previous studies have often been limited in scope and methodology due to their focus on broad age ranges and reliance on pairwise interactions based on pre-defined brain atlases to explore lifespan trajectories. Additionally, while data-driven ICA has been employed in lifespan studies, these studies have not examined multiscale interactions. As a result, these studies did not provide a comprehensive and precise view of developmental trajectories during the crucial first six months of infant brain development. Additionally, the role of multiway network interactions in connectome growth during this critical period has yet to be studied, as pairwise interactions may overlook significant high-order functional connections in the brain [29, 30, 31, 33, 34, 39].

To address these critical knowledge and technique gaps, this study explored the continuous developmental trajectory of multiway, multiscale network interactions over the first 180 days and examined its relationship with infant age. Firstly, we utilized high-quality longitudinal neuroimaging data, including 126 scans from 71 typically developing infants. These infants underwent resting-state functional MRI scans at various ages, from birth to six months. Secondly, we utilized multi-scale intrinsic connectivity networks (ICNs) derived from the infants in our study, building on the foundational work of a recent study that identified 105 multi-scale ICNs in infants aged 0 to 6 months [40]. This prior study employed both blind data-driven and reference-informed methods to establish the existence of these ICNs during infancy. The identified ICNs align with the *NeuroMark 2*.*1 template*, a universal model derived from over 100K subjects, offering networks resolved across multiple spatial scales [33].

To comprehensively map the developmental trajectory and capture higher-order functional connections, we assessed both pairwise and triple interactions, along with their respective subspaces, and examined their correlation with infant age to identify key networks. Including both interaction types is crucial, as pairwise interactions alone may not fully capture the complexity of brain connectivity, particularly during early development when networks are rapidly evolving. By investigating both pairwise and triple interactions and their subspace, we are able to reveal a more complete and nuanced understanding of the brain’s functional architecture. This dual approach enables us to identify not only direct interactions between brain networks but also the higher-order relationships that are critical for understanding the complex dynamics of early brain development.

## 2 Results

### 2.1 Estimated Subject-Specific ICNs and Their Time Courses Using Infant rsfMRI

To estimate subject-specific ICNs and their time courses, we applied the spatially constrained ICA method, Multivariate-Objective Optimization (MOO-ICAR) [33, 41], using the *NeuroMark2*.*1 Template* as a reference to obtain subject-specific multi-scale ICN spatial maps and their time courses in infants. Finally, we categorized 105 multi-scale ICNs into the following domains: visual (VI, 12 ICNs, includes visual network, VIS; ventral attention network, VAN), cerebellar (CB, 13 ICNs), temporal (TP/TMN, 13 ICNs, includes paralimbic network, PLN; explicit memory network, EM), subcortical (SC, 23 ICNs, includes limbic network, LIM; explicit memory network, EM; language network, LAN), somatomotor (SM, 13 ICNs), and higher cognitive (HC, with 31 ICNs, includes the default mode network, DMN; language network, LAN; central executive network, CEN; salience network, SN; and dorsal attention network, DAN), as shown in **Fig.1A**.

**Figure 1:**
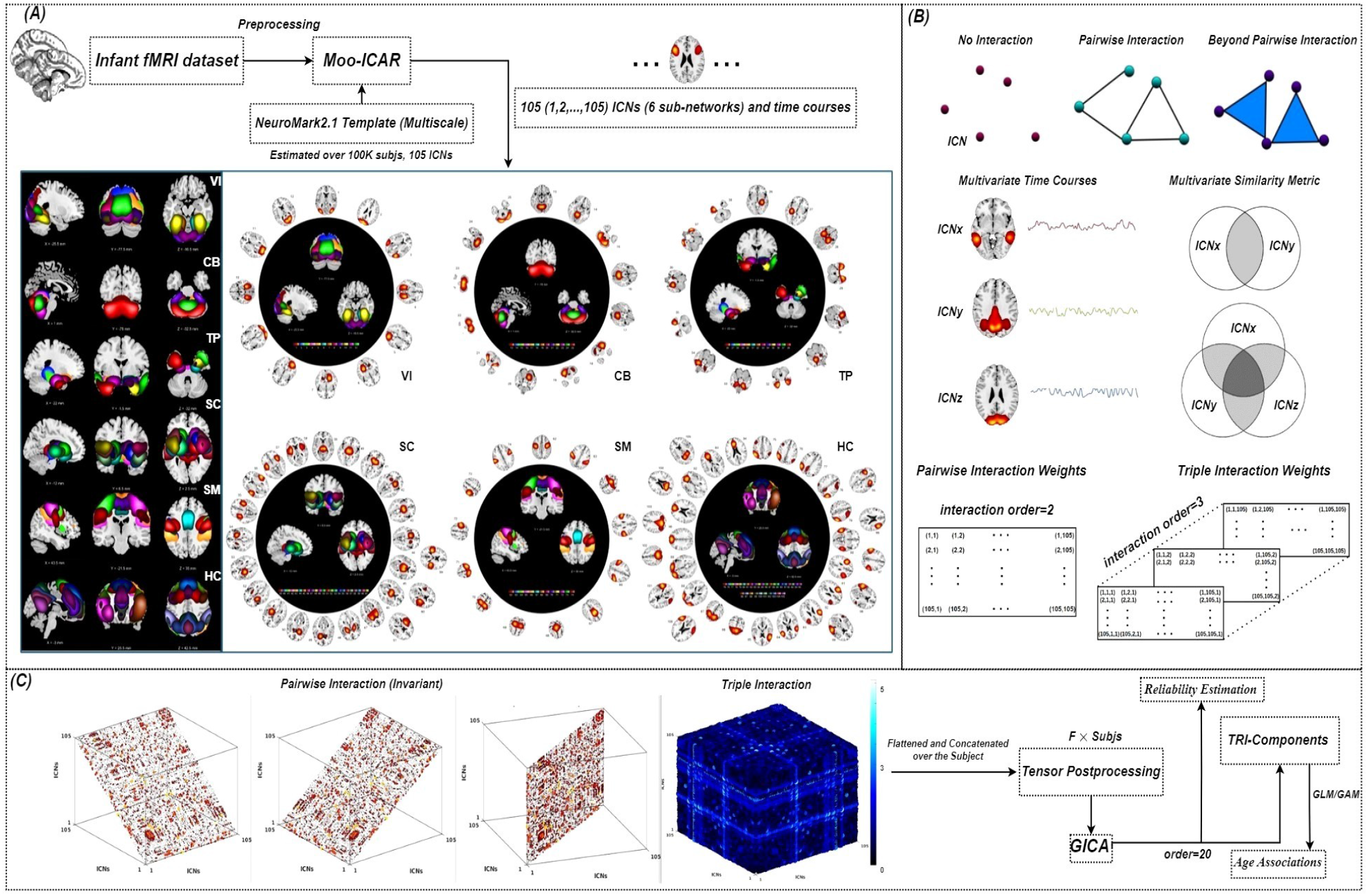
Framework for analyzing multiway multiscale interactions within brain networks. The preprocessed infant resting-state fMRI data were input into Multivariate Objective Optimization ICA with Reference (MOO-ICAR) using the spatially constrained *NeuroMark2*.*1 Template*, derived from over 100K subjects [33]. This process yielded subject-specific estimates of 105 intrinsic connectivity networks (ICNs) and their corresponding time courses in infant rsfMRI. These 105 ICNs were then grouped into six brain domains: visual (VI), cerebellar (CB), temporal (TP/TMN), subcortical (SC), somatomotor (SM), and higher cognitive (HC), as depicted in ***A***. As we know, the network in the human brain is not isolated; interactions occur both pairwise and beyond pairwise for information exchange. In this study, we estimated pairwise (interaction order = 2) and triple interactions (interaction order = 3) among intrinsic connectivity networks (ICNs), shedding light on multiway interactions in the infant brain, as shown in ***B***. We first estimated the pairwise interactions for each subject and then concatenated these pairwise interactions (105× 105) across subjects (*F Subjs*, where *F* represents the total number of features). For triple interactions, we flattened the 3D interaction tensors (105× 105 × 105) after estimating them and then concatenated them across subjects. Due to the high dimensionality of triple interactions and to explore the latent space underlying these complex pairwise and triple interactions, we applied group independent component analysis (GICA) to decompose the intricate patterns of both pairwise and triple interaction components. Subsequent analyses then focused on exploring the relationship between age and the decomposed brain network interaction components, as illustrated in ***C***.

### 2.2 Estimated Triple and Pairwise Interactions from Infant rsfMRI

Here, we estimated triple and pairwise interactions between specific ICNs using total correlation (for interaction order 3, i.e., *TC*(*ICN*_*x*_, *ICN*_*y*_, *ICN*_*z*_)), mutual information (for interaction order 2, i.e., *MI*(*ICN*_*x*_, *ICN*_*y*_)), and Pearson correlation (i.e., *PC*(*ICN*_*x*_, *ICN*_*y*_)), respectively, as illustrated in **Fig.1B**. In total, there are 105^3^ possible triple interactions when considering all combinations, including 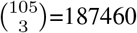 unique sets of triple interactions. Similarly, there are 105^2^ possible pairwise interactions, which include 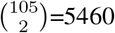 unique sets of pairwise interactions. The estimated pairwise interactions for each subject are represented as 2D matrices, while triple interactions are represented as 3D tensors, as shown in **Fig.1C**.

### 2.3 Triple and Pairwise Functional Connectivity in the Infant Brain and Their Associations with Age

We investigate the relationship between brain development and age by examining the association of the triple and pairwise functional connectivity with age. The correlation between age and z-scored functional connectivity for both triple and pairwise interactions is shown on the left side of **Fig.2A**,**B**. Additionally, networks with triple and pairwise interactions showing significant associations with age were identified, as presented on the right side of **Fig.2A**,**B**. We observed a cluster pattern along the diagonal in triple functional connectivity, while a similar pattern appeared in pairwise functional connectivity. Additionally, triple functional connectivity estimated more connections compared to pairwise connectivity, potentially offering deeper insights into brain functional connectivity.

**Figure 2:**
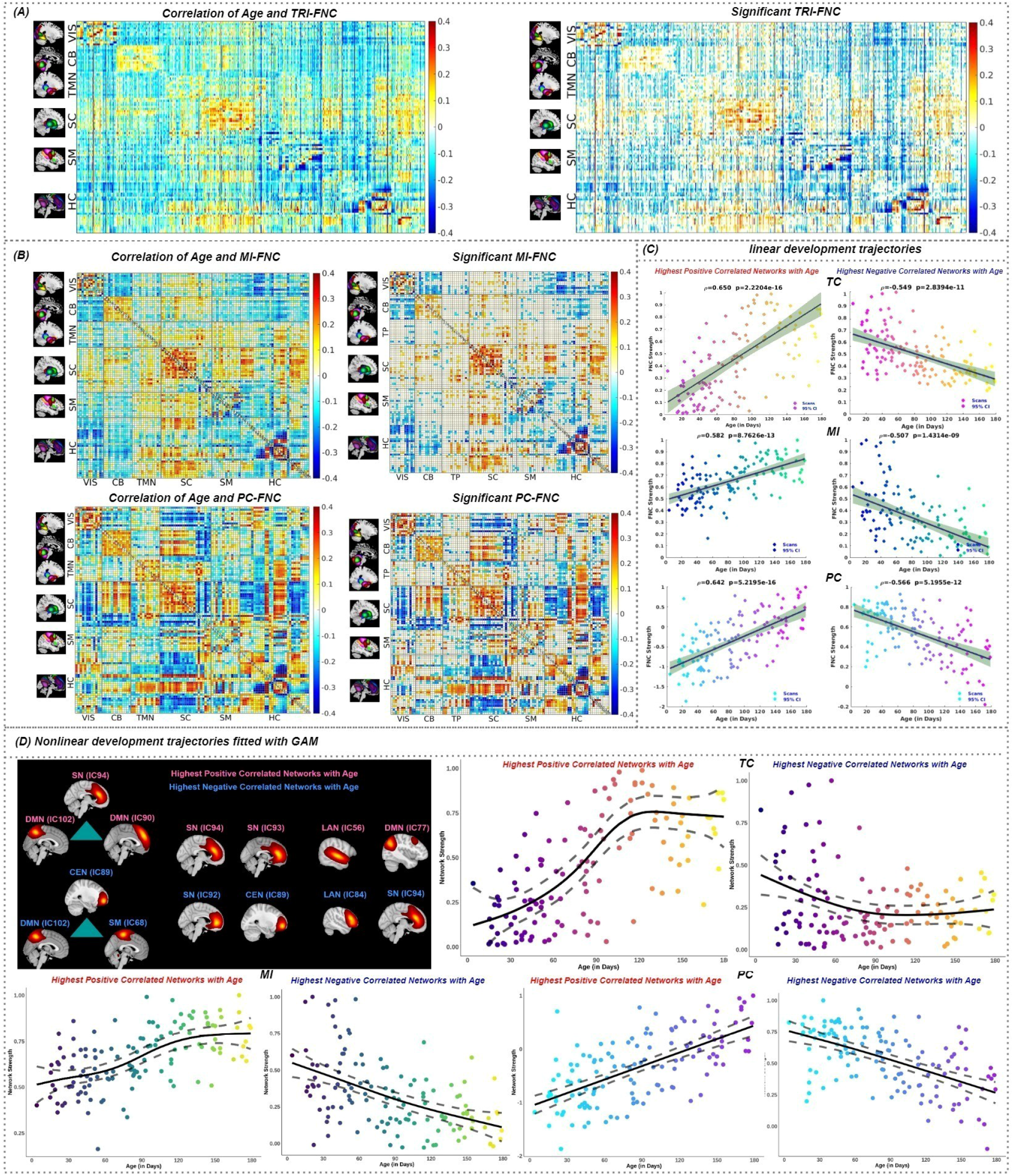
Triple and pairwise brain network interactions in the infant brain and their associations with age. In ***A***, the correlation between z-scored triple functional connectivity (TRI-FNC) and age is presented. Significant triple networks are identified on the right, with related domains labeled. A similar analysis was conducted for MI and PC, as shown in ***B***. To further evaluate network interactions with age, we identified the strongest positive and negative networks from both triple and pairwise functional connectivity. Their linear correlations are illustrated in ***C***. Additionally, ***D*** presents the nonlinear associations with age, estimated using generalized additive models (GAM). The Restricted Maximum Likelihood (REML) was employed within the model. This includes the strongest positive and negative triple interactions ([94, 102, 90] vs. [89, 102, 68]) and pairwise interactions ([94, 93] vs. [92, 89]; [56, 77] vs. [84, 94]).

Subsequently, we identified specific networks demonstrating the highest positive and negative correlations with age from both triple and pairwise interactions. The networks showing the highest positive correlation with age in triple interactions were SN (IC94), DMN (IC102), and DMN (IC90). The plot of network interaction values across age, displayed on the left side of **Fig.2C**, reveals a clear upward trend. The correlation value for this trend was calculated at 0.650, *p <* 0.001 (Bonferroni corrected), indicating a strong positive association between these network interaction values and age.

Conversely, the networks exhibiting the highest negative correlation with age in triple interactions were CEN (IC89), DMN (IC102), and SM (IC68). The fitted linear line, shown on the right side of **Fig.2C**, demonstrates a downward trajectory, suggesting a decreasing trend with age. The correlation value for this trend was computed at -0.549, *p <* 0.001 (Bonferroni corrected), reflecting a substantial negative correlation between these network interaction values and age.

The same analysis was performed for pairwise interactions using MI and PC, respectively. For pairwise interactions, the networks demonstrating the highest positive correlation with age were between the SN (IC94) and SN (IC93), and between the LAN (IC77) and DMN (IC56). The plot of network interaction values across age, displayed on the left side of **Fig.2C**, shows an upward trend. The correlation values for this trend were calculated at 0.582 for MI and 0.642 for PC, with *p <* 0.001 (Bonferroni corrected), indicating a strong positive association between these network interactions and age.

Similarly, the networks showing the highest negative correlation with age in pairwise interactions were between SN (IC92) and CEN (IC89), and between LAN (IC84) and SN (IC94). The fitted line, shown on the right side of **Fig.2C**, demonstrates a downward trajectory, suggesting a decreasing trend with age. The correlation values for this trend were computed at -0.507 for MI and -0.566 for PC, with *p <* 0.001 (Bonferroni corrected), indicating a substantial negative correlation between these network interactions and age.

To capture nonlinear patterns in network interactions with age, we employed a generalized additive model (GAM) to fit network interactions and age. We observed distinct S-shaped nonlinear trajectories, with the strongest patterns evident in positive correlation triples (TC, SN IC94, DMN IC102, DMN IC90) and MI (SN IC94, SN IC93) interaction networks compared to PC (LAN IC77, DMN IC56) interactions. This suggests that brain development experiences a sharp developmental period in the early stages and then tends to stabilize as it consolidates these shaped connections. Conversely, we observed that interactions in the negative correlation network tend to show a decreasing shape with age. Moreover, as shown in **Fig.2D**, interaction estimates from TC and MI exhibit similar nonlinear S-curve shapes, unlike those from PC. This similarity arises because TC and MI capture nonlinear dependencies, whereas PC does not, which likely explains the observed differences.

### 2.4 Triple Interactions Capture Different Brain Development Information than Pairwise Interactions

The information about brain development derived from triple interactions differs from that obtained through pair-wise interactions alone, as illustrated in **Fig.3A**. To explore these differences, we analyzed the highest positive and negative triple correlation networks with age by breaking them down into pairwise interactions. First, we observed that triple interaction networks show substantial positive and negative correlations with age (*ρ*=0.650 vs. *ρ*=-0.549, p<0.001, Bonferroni corrected), as presented in **Fig.3B**. We decomposed the triple interaction into pairwise interactions and then evaluated the average combined pairwise interactions 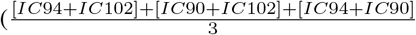 vs. 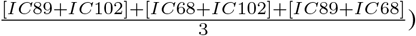 for MI and PC separately. We observed that the average combined pairwise interactions generally exhibit weak negative associations with age (*ρ*=-0.067 vs. *ρ*=-0.186 for MI, and *ρ*=-0.276 vs. *ρ*=-0.127 for PC). These results, as illustrated on the left side of **Fig.3C**, highlight the differing patterns between the two measures. Meanwhile, we also examined pairwise interactions (IC94 vs. IC102; IC94 vs. IC90; IC102 vs. IC90 in the highest positive triple interaction networks, and IC89 vs. IC102; IC89 vs. IC68; IC102 vs. IC68 in the highest negative triple interaction networks) separately using MI and found similar results (*ρ*=-0.125 vs. *ρ*=-0.107; *ρ*=0.072 vs. *ρ*=-0.051; *ρ*=-0.130 vs. *ρ*=-0.315). In summary, we demonstrated that triple interactions capture different brain development network interactions than pairwise interactions, and that pairwise interactions alone may miss some information that is captured in triple interactions.

**Figure 3:**
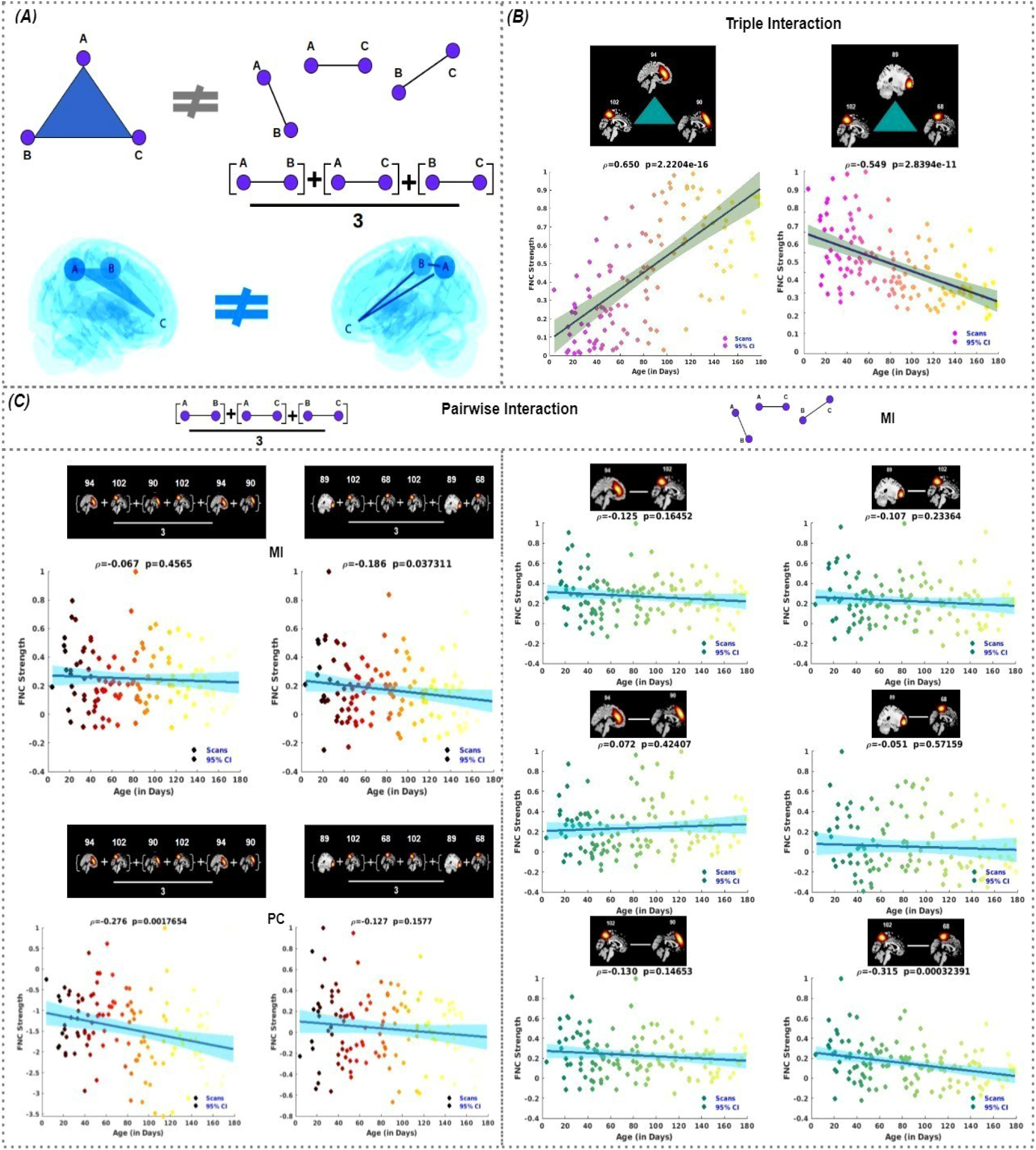
Triple interactions reveal stronger associations with age compared to pairwise interactions. The triple interaction (e.g., [A, B, C]) can be broken down into pairwise interactions ([A, B], [A, C], [B, C]) or their combined average. However, these decomposing pairwise interactions are not equivalent to the original triple interaction, as illustrated in ***A***. To further illustrate this distinction, we examined how these interactions relate to brain development. ***B*** shows the strongest positive and negative triple network interactions across different infant ages. In ***C***, we compare these interactions by presenting the associations derived from both the combined average of pairwise interactions and purely pairwise interactions, which are extracted from the z-score pairwise functional connectivity matrix. This comparison highlights the additional brain development information provided by triple interactions that is not captured by pairwise interactions alone.

### 2.5 Identified Latent Network Connectivity Subspaces in Triple and Pairwise Interactions from ICA

By employing ICA with a model order of 20, selected based on the elbow criterion, we successfully separated the complex pairwise and triple interaction patterns into 20 distinct components, as illustrated in **Fig.4A**. This decomposition is essential for gaining a deeper understanding of the independent structures within infant brain networks. Examining both the individual components and the interactions themselves allows us to explore their contributions to brain function. Moreover, this approach enables us to extract latent space information, enriching our analysis of the dynamic connectivity in infant brains.

**Figure 4:**
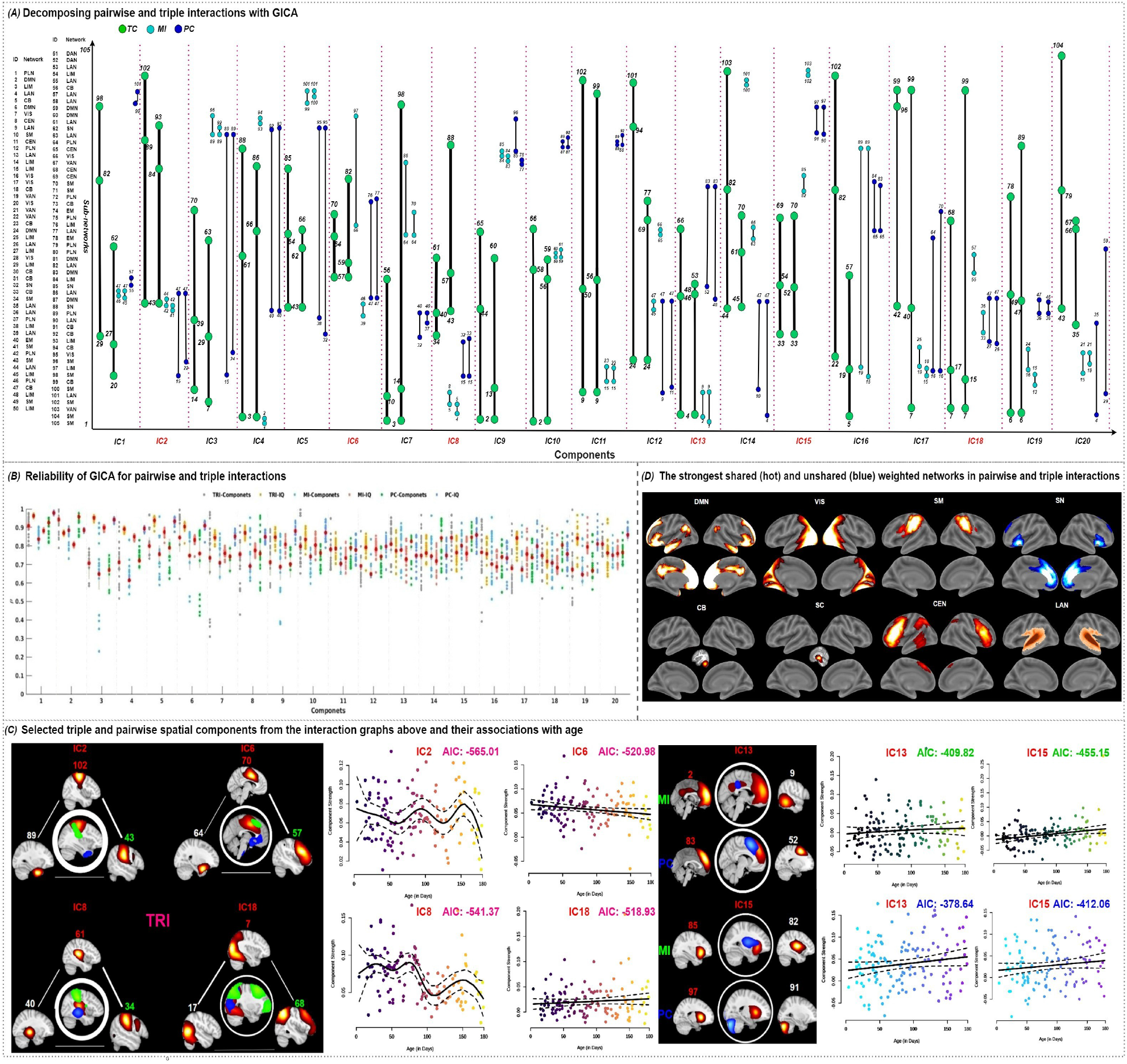
Decomposed latent triple and pairwise network connectivity subspaces in the infant brain and their associations with age. In panel ***A***, the triple and pairwise subspace latent interactions are identified, with the first and second strongest interactions for each component displayed from left to right. Interactions estimated from TC, MI, and PC are depicted in green, cyan, and blue, respectively. The indices correspond to specific ICNs listed on the left. The reliability of ICA for both pairwise and triple interactions was assessed by analyzing spatial similarities and stability values (IQ) from 20 triple components (TRI-C) obtained through ICA after 20 repetitions. For comparison, similar analyses were performed for pairwise interactions using mutual information and Pearson correlation. The resulting similarities and corresponding IQ values for these 20 components are also presented, as shown in panel ***B***. Furthermore, the strongest triple interactions were identified in components 2, 6, and 8, while the weakest interactions were found in component 18. Components 13 and 15 were selected for pairwise interactions, as shown in panel ***C***. Additionally, pairwise interactions within triple interactions are also presented, with circle and line thickness indicating interaction strength. The association between age and the selected triple and pairwise components is presented, along with the corresponding Akaike Information Criterion (AIC) values to quantify the fitting performance (a lower value means better performance). The strongest shared and distinct weighted brain domains are displayed in panel ***D***. The most shared domains across both interaction types include default mode (DMN), visual (VIS), somatomotor (SM), cerebellar (CB), subcortical (SC), and central executive domain(CEN). Notably, one domain (language, LAN) was detected by both TC and MI, and another domain (salience, SN) was detected exclusively through TC, but neither was observed in pairwise interactions.

To ensure that the components produced by ICA are stable and reliable, we evaluated the reliability of both triple (TRI-C) and pairwise (PA-C: PC and MI) independent components. We report the related spatial similarities for these components (TRI-C vs. PA-C) and their stability values (IQ) (TRI-IQ vs. PA-IQ) in **Fig.4B**. The results indicate that all triple and pairwise components exhibit high spatial similarities across the 20 independent components, demonstrating stable reliability of ICA, with triple components generally performing better than pairwise components.

### 2.6 Latent Triple and Pairwise Network Connectivity Subspaces in the Infant Brain and Their Associations with Age

The ICA resulted in the decomposition of both the triple and pairwise interaction tensors into 20 distinct latent triple (TRI-C) and pairwise (PA-C) network connectivity subspaces, respectively. These components represent the most independent patterns of triple and pairwise interactions observed in the infant brain. The isolated 20 components for both triple and pairwise interactions were visually represented in the graph shown in **Fig.4A**, and we observed that hierarchical multiway network interactions occurred during infant development, involving multiple brain networks. The separate components of triple and pairwise interaction are illustrated in **Appendices Fig.1 and Fig.4**, respectively.

To thoroughly investigate the decomposed latent TRI-C and PA-C network connectivity subspaces in infant brain development, we analyzed several of the strongest and weakest components, focusing on their distinct patterns and how these patterns relate to developmental trajectories across different infant ages. Specifically, we focused on TRI-C2, TRI-C6, TRI-C8, and TRI-C13, which represent the strongest triple interactions, and TRI-C18, which represents the weakest triple interaction. Additionally, we analyzed PA-C13 and PA-C15 to identify the most significant and weakest pairwise interactions. All TRI-Cs and PA-Cs spatial maps are presented in **Appendices Fig.2 and Fig.5**, respectively.

**Figure 5:**
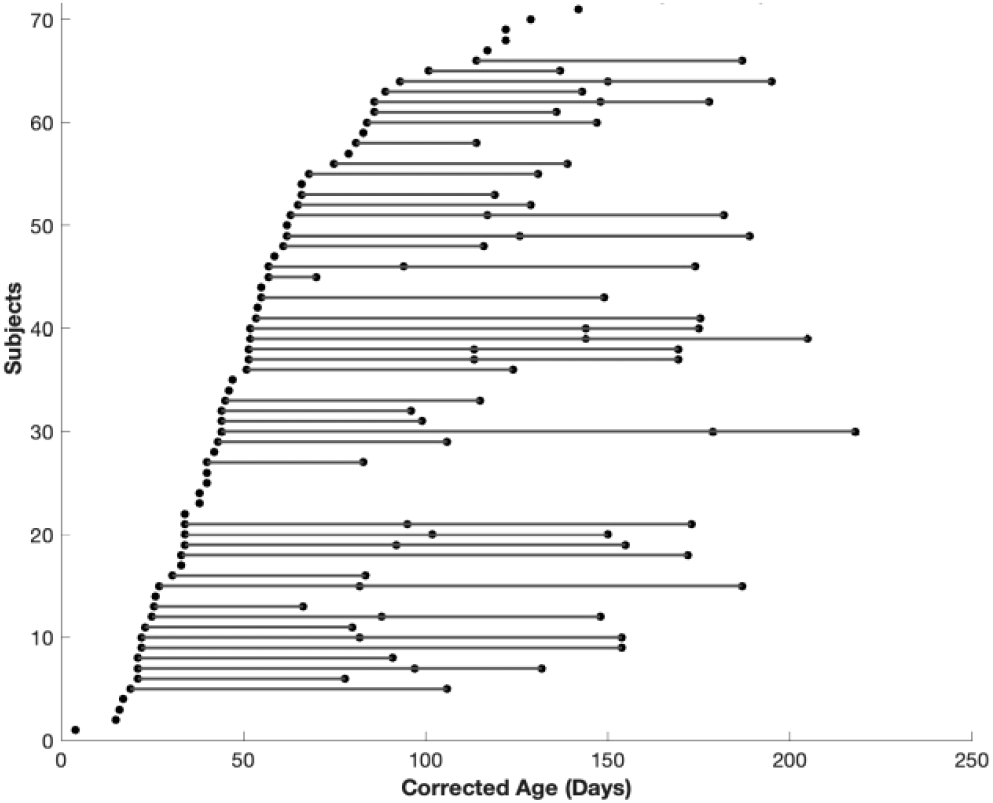
Distribution of age at scan among all participants. Each dot represents a single scan of a participant and the longitudinal scans from the same participant are connected by lines.

In the analysis of the strongest triple interactions in TRI-C2, we examined both the components and interactions to gain a comprehensive understanding of early brain development. The components reveal independent connectivity patterns, while the interactions show how these patterns combine to support brain function, capturing the complexity of brain network organization. Our analysis identified that the ICNs with the highest contributions—ICN102, ICN43, and ICN89—correspond to the SM, SM, and TP domains, respectively (see **Fig.4C**). This suggests that the strongest interactions within this component are primarily driven by the SM and TP domains, with the other ICNs contributing less to the component. Furthermore, we noted stronger pairwise interactions between ICN102 and ICN43 compared to other pairwise interactions, likely due to both ICs being part of the SM domains, where intra-network interactions are generally more robust than inter-network interactions.

In TRI-C6, we observed that the LAN (IC57) is involved in triple interactions with the SM (IC70) and LIM (IC54). Similar interactions are also observed in TRI-C8 (SM-LAN-EM) and TRI-C18 (VIS-VIS-CEN). Furthermore, in PA-C13, we identified the strongest pairwise interactions between the DMN (IC2 and IC83) and LAN (IC9) and DAN (IC52). Additionally, in PA-C15, the weakest interactions were observed between the LAN (IC82) and SN (IC85), as well as between the LAN and LIM (IC97) and CB (IC91).

To further investigate the relationship between decomposing TRI-Cs and PA-Cs and infant age, we applied a generalized additive model (GAM). The results of this analysis are presented in **Fig.4C**. Additionally, to quantify the model fit performance, we reported the Akaike Information Criterion (AIC) value, which indicated that TRI-Cs provide a better fit than PA-Cs. The relationships between all TRI-Cs and PA-Cs with infant age were presented in **Appendices Fig.3, Fig.6 and Fig.7**, respectively.

**Figure 6:**
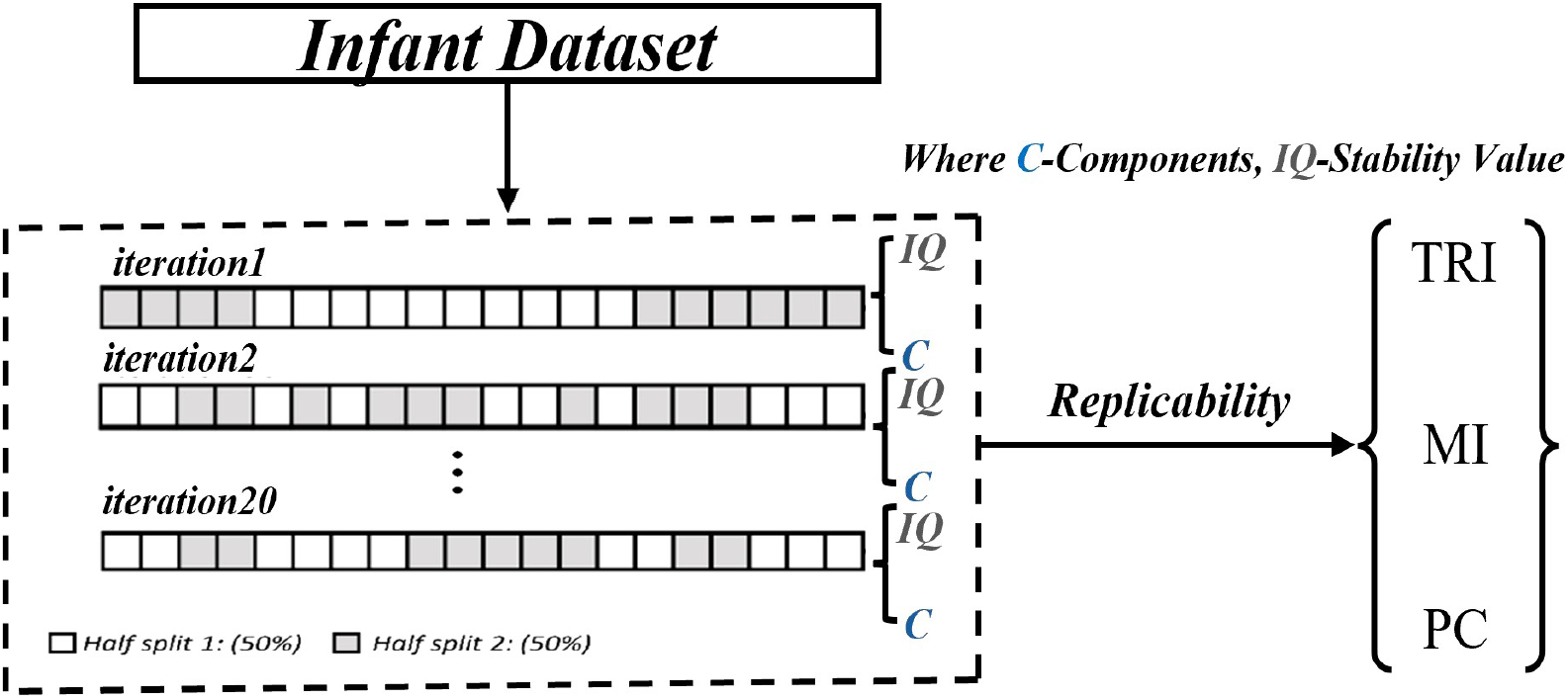
Pipeline to assess the reliability of the ICA approach. The entire infant dataset was randomly split in half. ICA was applied to each half-split, concatenating two-way and triple interactions across subjects to generate 20 components and their corresponding IQ. This process was independently repeated 20 times. Finally, the average spatial similarity was calculated.

Based on the hierarchical triple and pairwise interactions identified in **Fig.4A**, we observed the engagement of multiple ICNs. To determine which brain domains significantly contribute to infant brain development, we identified several commonly shared brain domains across both triple (by TC) and pairwise interactions (by MI and PC), including the DMN, VIS, SM, CB, LIM, and CEN, as shown in **Fig.4D**. This analysis indicates that most of these domains emerge within the first six months of life. Additionally, we noted that the LAN was detected only by TC and MI, but not by PC. One major reason for this is that PC may not capture some nonlinear interactions effectively. Furthermore, MI was also unable to detect the SN compared to TC, as illustrated in **Fig.4D**. In summary, triple interactions reveal certain domains in infants that are not captured by pairwise interactions, highlighting the added value of analyzing hierarchical triple interactions for a more comprehensive understanding of early brain development.

## 3 Discussion

Investigating multiway multiscale interactions in extensive brain networks during the first 180 days of life provides crucial insights into early brain development and potential early markers for neurodevelopmental disorders. This research, which includes 71 infants within their first 180 days, employs spatially constrained ICA to derive multiscale infant brain networks and then examines multiway interactions within these networks. Pairwise and triple interactions are estimated using PC, MI, and triple interaction models, with their associations to infant age assessed. Additionally, the subspace latent triple and pairwise interactions and their associations with age are explored. The findings from this work are significant, as they lay the groundwork for a deeper understanding of the brain’s early functional organization and its developmental trajectory.

Our studies revealed that multiway network interactions occur during infant brain development. These brain domains encompass high-level cognitive processes as well as visual, attention, and language networks. In our study, we also found several key domains captured by all metrics in the infant brain during the first six months, including the DMN, VIS, SM, CB, LIM, and CEN. Additionally, we discovered that one domain, LAN, was detectable through TC and MI but not through PC. This underscores the importance of considering nonlinearity when estimating network interactions. Furthermore, we found that another domain, SN, was detectable only through TC and not through MI or PC. This highlights the need to estimate high-order network interactions when studying brain function, as pairwise interactions alone do not capture these more complex relationships.

To understand how network interactions change with infant age, we first examined general patterns of age-related associations across different network domains. We then focused on the most significant interactions by identifying the highest positive and negative network interactions from both triple and pairwise analyses, and assessed their linear and nonlinear statistical relationships. The three most prominent domains identified with the highest positive and negative triple interactions were the DMN, CEN, and SN. Additionally, DMN, SN and LAN were also identified from pairwise interactions. Interestingly, these three networks also form a canonical triple network model [42, 43], and their irregular interactions are linked to various brain disorders, including autism spectrum disorders (ASD) [7, 44]. Therefore, these networks play a crucial role in early infant brain development, particularly within the specific six-month window. Furthermore, the distinct nonlinear trajectories observed in our analysis suggest a dynamic and complex pattern of brain development, particularly in the interaction networks associated with positive correlations, such as those between the TC, SN IC94, DMN IC102, and DMN IC90. These patterns indicate that certain brain networks undergo rapid, nonlinear changes in the early stages of development, followed by a phase of stabilization as the brain consolidates and solidifies these connections. This sharp developmental period in the initial stages could reflect critical phases of brain maturation, where fundamental networks like the SN and DMN form and refine their interactions. The contrast in the fitting shape with age observed in the interaction networks involving MI (such as SN IC94 and SN IC93) and PC (LAN IC77, DMN IC56) further supports the idea that brain regions associated with higher-order cognitive functions and attention networks may undergo more significant changes during early development.

The eventual stabilization of these trajectories aligns with current models of brain development, where the early brain is marked by heightened plasticity, allowing for rapid changes and refinements in functional connectivity [7, 8]. As the brain matures, these networks converge toward more stable configurations, reflecting a consolidation of functional pathways that support mature cognitive and behavioral processes. Our findings suggest that this developmental trajectory is not uniform across all brain networks.

To further elaborate on the differences between triple and pairwise interactions, we estimated pairwise interactions from the highest positive and negative triple interactions and evaluated their relationships in comparison to triple interactions. We investigated two approaches for analyzing pairwise interactions: one method involved examining each pairwise interaction individually, while the other involved averaging the combined pairwise interactions. We found that pairwise interactions alone significantly reduced the relationship between network interaction strengths and age compared to triple interactions. Moreover, after applying ICA to both triple and pairwise functional connectivity, we identified latent network connectivity subspaces within these interactions. We then explored their correlation with age, and our results also demonstrate that the triple interaction networks provide a better explanation for age-related changes than the pairwise interactions. This indicates that triple interactions provide richer information than pairwise interactions, which is valuable for capturing more comprehensive information to deepen our understanding of changes in brain network interactions during early infant brain development.

Taken together, the analysis of multiway interactions in infant brain connectivity, while offering valuable insights into intricate network interactions, is inherently limited by several factors that reflect the broader challenges of studying multi-order interactions in the infant brain.

### 3.1 Limitations

#### 3.1.1 Computational Complexity and Biological Interpretation

One major limitation is the computational complexity associated with triple interactions [22, 24, 29]. The complexity of analyzing triple interactions is markedly higher than that of pairwise interactions, leading to increased computational demands. As the number of possible interactions grows combinatorially with the number of ICNs, this can strain computational resources and extend analysis time. The interpretation of such interactions is inherently complex; unraveling the contributions of each individual component and understanding their joint effects on infant brain function is often challenging. Unlike pairwise interactions, which provide a relatively straightforward measure of connectivity between ICNs, triple interactions involve understanding how three ICNs influence each other simultaneously. This complexity can obscure the underlying mechanisms of brain function and make it difficult to pinpoint the exact nature of the interactions [29]. Moreover, the biological relevance of these interactions is not always clear [45]; Overall, while high-order interaction analyses hold promise for advancing our understanding of brain connectivity, balancing computational demands with the need for clear biological interpretation remains a key challenge. Additionally, triple interaction analyses are highly affected by data quality issues like noise and inaccuracies in preprocessing, which can significantly impact the results [46, 47, 45].

#### 3.1.2 Beyond Triple Interactions in the Human Brain

Another critical consideration is the nature of interactions beyond triple interactions [26, 29, 30, 45]. The brain’s connectivity is not limited to simple triplets but involves a complex interplay of interactions at multiple levels. Higher-order interactions, such as quadruple or quintuple interactions, and the mixing of different types of interactions (e.g., linear and nonlinear) further complicate the analytical landscape. The interplay between various interaction orders and their combined effects can create a tangled web of connectivity that is challenging to untangle and interpret. Furthermore, we will also face computing and memory challenges as we use higher-order interactions, which will require significant computational resources.

#### 3.1.3 Alternative Approaches for Estimating Multiway Interactions

In addition, in this study, we utilized TC to estimate triple interactions in infant brain connectivity, which provides valuable insights into the intricate dependencies among three ICNs. TC measures the overall dependence among three variables by quantifying how much the joint distribution deviates from the product of the marginal distributions. However, there are also other methods to estimate high-order interactions, such as hypergraphs [48], persistent diagrams [49], dual total correlation [50], and O/S information [51]. Each of these metrics offers a different approach to capturing high-order interactions, and exploring these methods in future research could provide additional valuable insights. Specifically, O/S information is noteworthy because it can quantify the balance between redundancy and synergy in the human brain, and it is calculated from TC and dual total correlation [51].

#### 3.1.4 Brain Activity: Dynamic Rather Than Static

In this study, we focused on analyzing triple interactions to understand connectivity patterns within the infant brain. However, it is essential to recognize that these interactions are inherently dynamic rather than static, which has significant implications for interpreting our findings and understanding brain connectivity [29, 32, 45, 52, 53]. By examining how interactions between infant brain regions change over time, dynamic functional connectivity methods can reveal transient states and fluctuations that are not apparent in static models [54, 55]. Therefore, integrating dynamic approaches into the study of high-order interactions is crucial for a more comprehensive understanding of brain connectivity. Additionally, exploring how these dynamic interactions relate to cognitive processes and behaviors will enhance our understanding of the functional network dynamics in the infant brain.

In summary, while analyzing triple interactions provides valuable insights into the development of infant brain connectivity beyond what is revealed by pairwise interactions, it also presents significant challenges that mirror the broader difficulties of studying complex multi-order interactions. Overcoming these challenges through rigorous statistical methods and exploring alternative approaches is essential for deepening our understanding of brain development and its impact on cognitive and behavioral growth.

## 4 Methods

### 4.1 Infancy Data

Infant scans were conducted at Emory University’s Center for Systems Imaging Core using either a 3T Siemens Tim Trio or a 3T Siemens Prisma scanner, each equipped with a 32-channel head coil [56, 57]. To ensure optimal conditions, all scans were performed while the infants were in a natural sleep state. We collected longitudinal resting-state functional magnetic resonance imaging (rsfMRI) data from 71 typically developing infants at up to 3 pseudorandom time points between birth and 6 months, resulting in a total of 126 scans, as depicted in Fig.5. The participants comprised 76 scans from 41 male subjects and 50 scans from 30 female infants aged from 4 to 179 days (mean corrected age ± Std. is 86.52 *±* 48.05 in days), covering the period from birth to approximately 6 months [12], as shown in Table.1. Informed written consent was obtained from the parents of all participants before their participation, and ethical approval for all study procedures was granted by Emory University.

**Table 1.** Participant characteristics at infancy, scan counts, subject counts, and corrected age (in days).

| | Scan Counts | Subject Counts | Mean Corrected Age ± Std. | Min. Corrected Age | Max. Corrected Age |
| --- | --- | --- | --- | --- | --- |
| Male | 76 | 41 | $90.58 \pm 48.66$ | 4 | 179 |
| Female | 50 | 30 | $80.35 \pm 46.92$ | 16 | 178 |
| Total | 126 | 71 | $86.52 \pm 48.05$ | 4 | 179 |

### 4.2 Data Acquisition

The fMRI data were acquired using two different scanner setups, both employing a multiband echo-planar imaging (EPI) sequence to enhance acquisition quality.

The fMRI was collected using a Siemens Tim Trio scanner. The acquisition parameters for this setup were a repetition time (TR) of 720 ms, an echo time (TE) of 33 ms, and a flip angle of 53^*°*^. The multiband factor was set to 6, with a field-of-view (FOV) of 208 ×208 mm and an image matrix of 84 ×84, resulting in a spatial resolution of 2.5 mm isotropic. To cover the entire brain, 48 axial slices were acquired. To improve the signal-to-noise ratio (SNR), 6 dummy scans were performed prior to the actual data acquisition to ensure steady-state magnetization.

The fMRI was acquired using a Siemens Prisma scanner. The parameters for this acquisition included a repetition time (TR) of 800 ms, an echo time (TE) of 37 ms, and a flip angle of 52^*°*^. The multiband factor was increased to 8, with the field-of-view (FOV) maintained at 208 ×208 mm and an image matrix of 104× 104, providing a spatial resolution of 2 mm isotropic. This setup included 72 axial slices to cover the entire brain. Similar to the previous setup, 6 dummy scans were conducted before the actual data acquisition to ensure steady-state and enhance the SNR.

### 4.3 Data Preprocessing

We performed standard NeuroMark preprocessing using the FMRIB Software Library (FSL v6.0, https://fsl.fmrib.ox.ac.uk/fsl/fslwiki/) and the Statistical Parametric Mapping (SPM12, http://www.fil.ion.ucl.ac.uk/spm/) toolboxes within the MATLAB 2020b environment. Initially, we discarded the first ten dummy scans showing significant signal changes. Subsequently, we corrected image distortions using field maps and SBRef data with reverse phase encoding blips, resulting in pairs of images with distortions in opposite directions. The applytopup tool was used to apply the field map coefficients and correct distortion in the fMRI data. After distortion correction, we performed slice timing to address slice-dependent delays using SPM. Following slice timing, we conducted head motion correction and realigned all scans to the reference scan.

Given the unique characteristics of infant fMRI data compared to adults, we implemented a two-step normalization process to align the infant data to a standardized Montreal Neurological Institute (MNI) space. Initially, we acquired the UNC-BCP 4D Infant Brain Template [58] (https://www.nitrc.org/projects/uncbcp_4d_atlas/) and registered it to the adult MNI space. Subsequently, we normalized the infant data to the UBC-BCP MNI template, specifically selecting the template corresponding to 4 months of age, which represents the median age of our infant dataset. Finally, the normalized fMRI data underwent spatial smoothing using a Gaussian kernel with a full width at half maximum (FWHM) of 6 mm.

### 4.4 Theoretical Analysis

#### 4.4.1 Spatially Constrained ICA on Infancy rsfMRI

A spatially constrained ICA (scICA) method known as Multivariate Objective Optimization ICA with Reference (MOO-ICAR) was implemented using the GIFT software toolbox (http://trendscenter.org/software/gift) [53]. The MOO-ICAR framework estimates subject-level independent components (ICs) using existing network templates as spatial guides [32, 36, 52, 53, 59, 60]. Its main advantage is ensuring consistent correspondence between estimated ICs across subjects. Additionally, the scICA framework allows for customization of the network template used as a spatial reference in the ICA decomposition. This flexibility enables either disease-specific network analyses or more generalized assessments of well-established functional networks suitable for diverse populations [33, 34, 36, 61, 62, 63].

The MOO-ICAR algorithm, which implements scICA, optimizes two objective functions: one to enhance the overall independence of the networks and another to improve the alignment of each subject-specific network with its corresponding template [36]. Both objective functions, 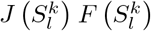,are listed in the following equation, which summarizes how the *l*^*th*^ network can be estimated for the *k*^*th*^ subject using the network template *S*_*l*_ as guidance:

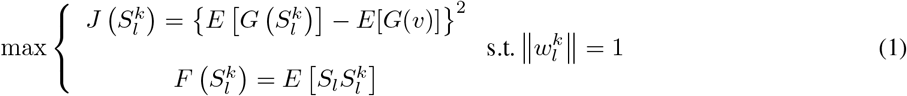

Here 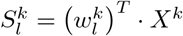 represents the estimated *l*^*th*^ network of the *k*^*th*^, *X*^*k*^ is the whitened fMRI data matrix of the *k*^*th*^ subject and 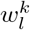 is the unmixing column vector, to be solved in the optimization functions. The function 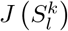 serves to optimize the independence of 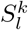 via negentropy. The *v* is a Gaussian variable with mean zero and unit variance *G*(.) is a nonquadratic function, and *E*[.] denotes the expectation of the variable. The function 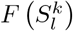 serves to optimize the correspondence between the template network *S*_*l*_ and subject network 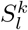.The optimization problem is addressed by combining the two objective functions through a linear weighted sum, with each weight set to 0.5. Using scICA with MOO-ICAR on each scan yields subject-specific ICNs for each of the N network templates, along with their associated time courses.

In this study, we used the *NeuroMark_fMRI_2*.*1 template* (available for download at https://trendscenter.org/data/) along with the MOO-ICAR framework for scICA on infant rsfMRI data. It enabled us to extract subject-specific ICNs and their associated time courses. This template includes N = 105 high-fidelity ICNs identified and reliably replicated across datasets with over 100K subjects [33]. These ICNs are organized into six major functional domains: visual domain (VI, 12 sub-networks), cerebellar domain (CB, 13 sub-networks), temporal-parietal domain (TP, 13 sub-networks), sub-cortical domain (SC, 23 sub-networks), sensorimotor domain (SM, 13 sub-networks), and high-level cognitive domain (HC, 31 sub-networks).

#### 4.4.2 Estimating Two-Way and Multiway Interactions in Infancy rsfMRI

##### Two-Way Interactions (Pearson Correlation)

The infancy rsfMRI signal comprising *n* ICNs obtained from scICA, denoted as *X*_*i*_ (1 ≤*i* ≤*n* and *n* = 105), each *X*_*i*_ corresponds informally to a time instance *t*_*i*_ in a sequence {*t*_1_, *t*_2_, …, *t*_*T*_} with a constant time interval *t*. Therefore, the two-way functional connectivity based on Pearson correlation can be estimated as follows:

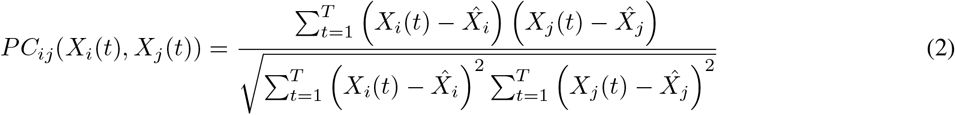

where 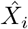 and 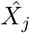 are the average of the rsfMRI signals at ICNs *i* and *j*. When two ICNs share substantial information, they are expected to exhibit a strong correlation, and conversely. In total, there are 105^2^ pairwise interactions based on Pearson correlation.

##### Two-Way Interactions (Mutual Information)

The mutual information between one ICN *X*_*i*_ and another ICN *X*_*j*_ is given by [64].

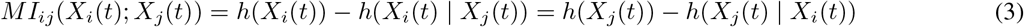

where the Shannon entropy *h*(*X*_*i*_(*t*)), *h*(*X*_*j*_(*t*)) of a continuous random variable *X*∈ *χ*with probability density function *f*_*X*_, its differential entropy is defined as [64],

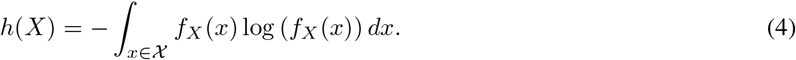

The mutual information measures the dependency between *X*_*i*_(*t*) and *X*_*j*_(*t*), and attains its minimum, equal to zero, if two ICNs independent. In total, there are 105^2^ pairwise interactions based on mutual information.

Although two-way interactions are commonly used to estimate functional connectivity in fMRI studies, we know that information interactions in the human brain usually go beyond two-way interactions. Considering the human brain as a high-dimensional and nonlinear complex system, relying solely on two-way interactions to explain brain function is insufficient and often overlooks higher-order interactions. Therefore, considering the limitations of two-way interactions [20, 24, 26, 29, 30], we applied total correlation to overcome these limitations and estimate higher-order information interactions in the human brain.

##### Multiway Interactions (Total Correlation)

The Total Correlation (TC) describe the dependence among *n* variables (*X*^1^, *· · ·, X*^*n*^) and can be considered as a non-negative generalization of the concept of mutual information from two parties to *n* parties. Let the definition of total correlation due Watanabe [28] be denoted as:

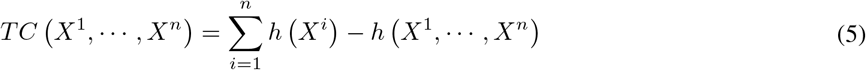

Where *h* (*X*^*i*^) is marginal entropy, and *h (X*^1^,, *X*^*n*^) is joint entropy. The TC will be equal to mutual information if we only have two variables, and TC will be zero if all variables are totally independent.

In real situations, estimating the marginal entropy *h* (*X*^*i*^) is straightforward, but estimating the joint entropy **h** (*X*^1^, *X*^2^*· · ·, X*^*n*^ *h* (*X*^*i*^) is considerably challenging. To address this challenge, Gaussian information theory is commonly applied to estimate total correlation because the BOLD signals satisfy Gaussian distributions [24, 31, 65, 66]. For a univariate Gaussian random variable *X* ∼ *𝒩* (*µ, σ*), the entropy (given in nats) will be,

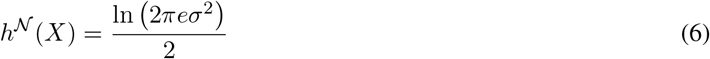

And for a multivariate Gaussian distribution, the joint entropy will be,

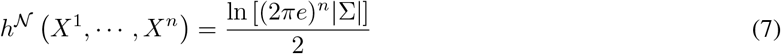

where refers to the determinant of the covariance matrix of ^1 *n*^. For the multivariate Gaussian case, the mutual information between (*X*^1^, *· · ·, X*^*n*^) and (*Y* ^1^, *· · ·, Y* ^*n*^) is given by,

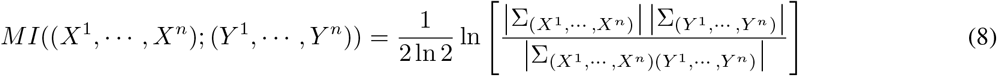

Finally, the Gaussian estimator for TC can be:

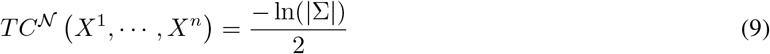

Now, we can estimate interactions beyond two-way (***k***=2) by computing ***k***-way interactions (***k***>2) among ***n*** = 105 ICNs, iterating over each set of indices used to obtain TC. In total, there are 105^3^ triple interactions if ***k*** = 3 is considered for all possible triple interactions, and there are 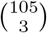 unique sets of triple interactions for multiscale infancy brain networks.

#### 4.4.3 Decomposing Two-way and Triple Interaction Tensors

Two-way interactions using PC and MI, as well as triple interaction tensors based on TC, were computed individually for each subject across 105 ICNs. Regarding triple interactions, we first extracted unique pairwise and triple interactions, flattened them, and then combined them across all subjects. These interactions were then aggregated across subjects and underwent blind ICA to extract the most independent components. To ensure the robustness and reliability of the ICA results, we executed the analysis 100 times using the Infomax optimization algorithm [67]. Each run included random initialization and bootstrapping techniques to mitigate potential biases and enhance the generalizability of the findings. This rigorous approach aimed to validate the stability and consistency of the identified components across multiple iterations, thereby reinforcing the overall validity of the ICA results [68, 69].

The decomposed components from the two-way and triple interactions were then ranked in descending order based on their self-weights. We focused on the top 10 weights and determined their corresponding indices to highlight the most distinctive triple-brain network interactions. We proceeded by evaluating the strength of these interactions across each ICN. Further analysis led us to identify the two strongest networks based on their aggregated weights. Subsequently, we pinpointed the networks that contributed most significantly to infancy development by examining interactions between the 105 ICNs and these top two networks.

#### 4.4.4 Assessing the Reliability of ICA for Two-Way and Triple Interactions in Infants

To measure reliability, we computed the average spatial similarity between corresponding ICNs from the two independent half-splits using Pearson correlation, as illustrated in Fig.6. Assessing the reliability of ICA for concatenated two-way (PC and MI) and triple (TRI) interactions across subjects is crucial for evaluating the stability and robustness of ICA [61]. In this study, we assessed the reliability of ICA for two-way and triple interactions by initially dividing the dataset randomly into two distinct subsets. Each subset underwent independent ICA processing, repeated 20 times. Each iteration produced components and their respective stability values (IQ), and the average spatial similarities were reported to evaluate the performance of ICA.

#### 4.4.5 Analyzing Linear and Nonlinear Association with Infant Age Using the General Linear Models and Generalized Additive Models

To investigate age-related changes in functional connectivity, we employed the general linear models (GLMs) [70] to assess the age effect on multiscale brain network interactions in infancy. We included gender, site and head motion parameters (mean FD) as covariates. The general linear model is defined as follows:

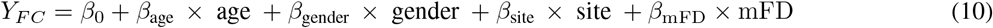

where *Y*_*F C*_ represents the connectome measures of each infant, *β*_age_ denotes the age effect. Bonferroni correction was applied to adjust for multiple comparisons across the measures, and significance was determined using a Bonferroni corrected *p <* 0.05.

To test the nonlinearity between brain networks and infant age, we applied generalized additive models (GAMs) [71]. GAMs are a flexible extension of generalized linear models (GLMs) that allow for nonlinear relationships between brain network interactions and infant age. In this context, GAMs are applied to model how functional connectivity changes with age, while also accounting for additional covariates such as gender, site, and motion. Mathematically, a GAM for modeling brain functional connectivity *Y*_*F C*_ can be expressed as:

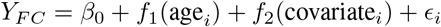

where *Y*_*F C*_ represents the functional connectivity measure for the *i*th ICN. *β*_0_ is the intercept term. *f*_1_(age_*i*_) is a smooth function of age, capturing the nonlinear relationship between age and functional connectivity. *f*_2_(covariate_*i*_) is a smooth function of other covariates (e.g., gender, site, and motion.). *ϵ*_*i*_ represents the residual error term, assumed to be normally distributed. The functions *f*_1_ and *f*_2_ are typically estimated nonparametrically using spline smoothing techniques, allowing for the flexible modeling of nonlinear effects. This approach enables the detection of complex patterns in functional connectivity that might be overlooked in traditional linear models, providing a more accurate understanding of how brain networks evolve over the lifespan. Here, we used Restricted Maximum Likelihood (REML) to select the optimal level of smoothing, which estimates smoothing parameters while accounting for the fixed effects. The method was implemented using the “mgcv” library in the R environment (https://cran.r-project.org/web/packages/mgcv/index.html). To quantify fitting performance, we calculated the Akaike Information Criterion (AIC) [72]. AIC assesses each model’s quality relative to others by balancing goodness of fit with model complexity, penalizing models with excessive complexity to prevent overfitting. Lower AIC values indicate a better balance between fit and complexity [72].

## 7 Ethics Statement

The infant datasets used in this work in under the study protocol approved by the Emory University Institutional Review Board (IRB). Legal guardians provided written consent for their infant’s participation at time of study enrollment.

## 8 Data and Code Availability Statement

Due to IRB restrictions, the infant data analyzed in this study cannot be shared without specific licenses. However, the dataset can be accessed upon request by contacting Dr.Sarah Shultz at *sarah*.*shultz@emory*.*edu*, who will facilitate the interaction with the IRB.

NeuroMark 2.1 templates are accessible on our lab website (https://trendscenter.org/data/) and GitHub (https://github.com/trendscenter/gift/tree/master/GroupICAT/icatb/icatb_templates). The UNC-BCP 4D Infant Brain Template is available at https://www.nitrc.org/projects/uncbcp_4d_atlas/. The codes of the GICA, and MOO-ICAR have been integrated into the group ICA Toolbox (GIFT 4.0c, https://trendscenter.org/software/gift/). The GAM model from the “mgcv” library in the R environment (https://cran.r-project.org/web/packages/mgcv/index.html).

## 9 Supporting Information

Additional supporting information can be found online in the Supporting Information section at the end of this article.

## 10 Appendices

Here, we present additional figures to support our claims in the results section.

Fig.1: TC interaction components from GICA.

Fig.2: TC spatial interaction components from GICA.

Fig.3: Age association with each TC interaction component from GICA.

Fig.4: MI&PC interaction components from GICA.

Fig.5: MI&PC spatial interaction components from GICA.

Fig.6: Age association with each MI interaction component from GICA.

Fig.7: Age association with each PC interaction component from GICA.

**Figure 1:**
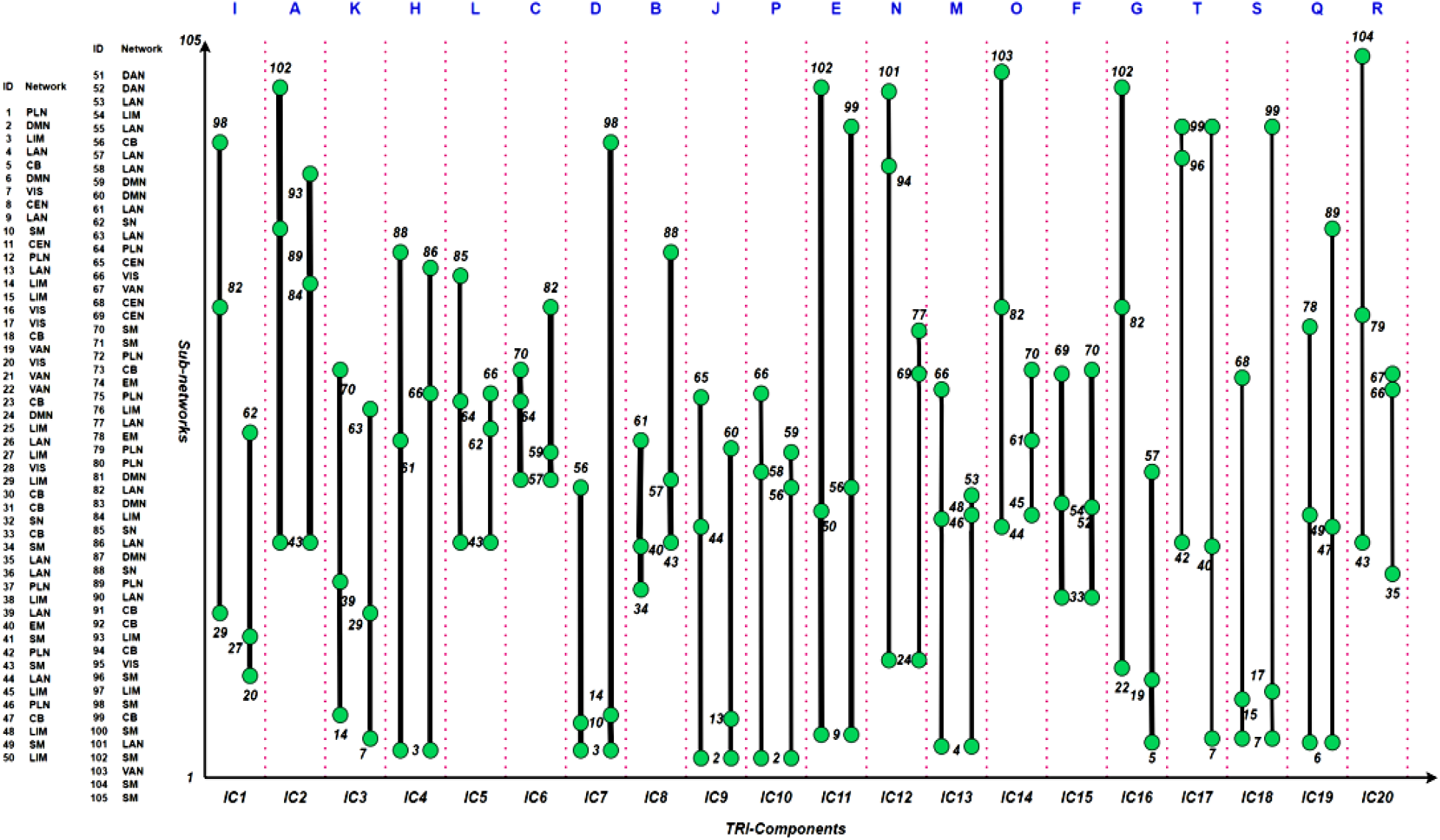
Triple interaction components from GICA. The order of the alphabet refers to network interaction strengths.

**Figure 2:**
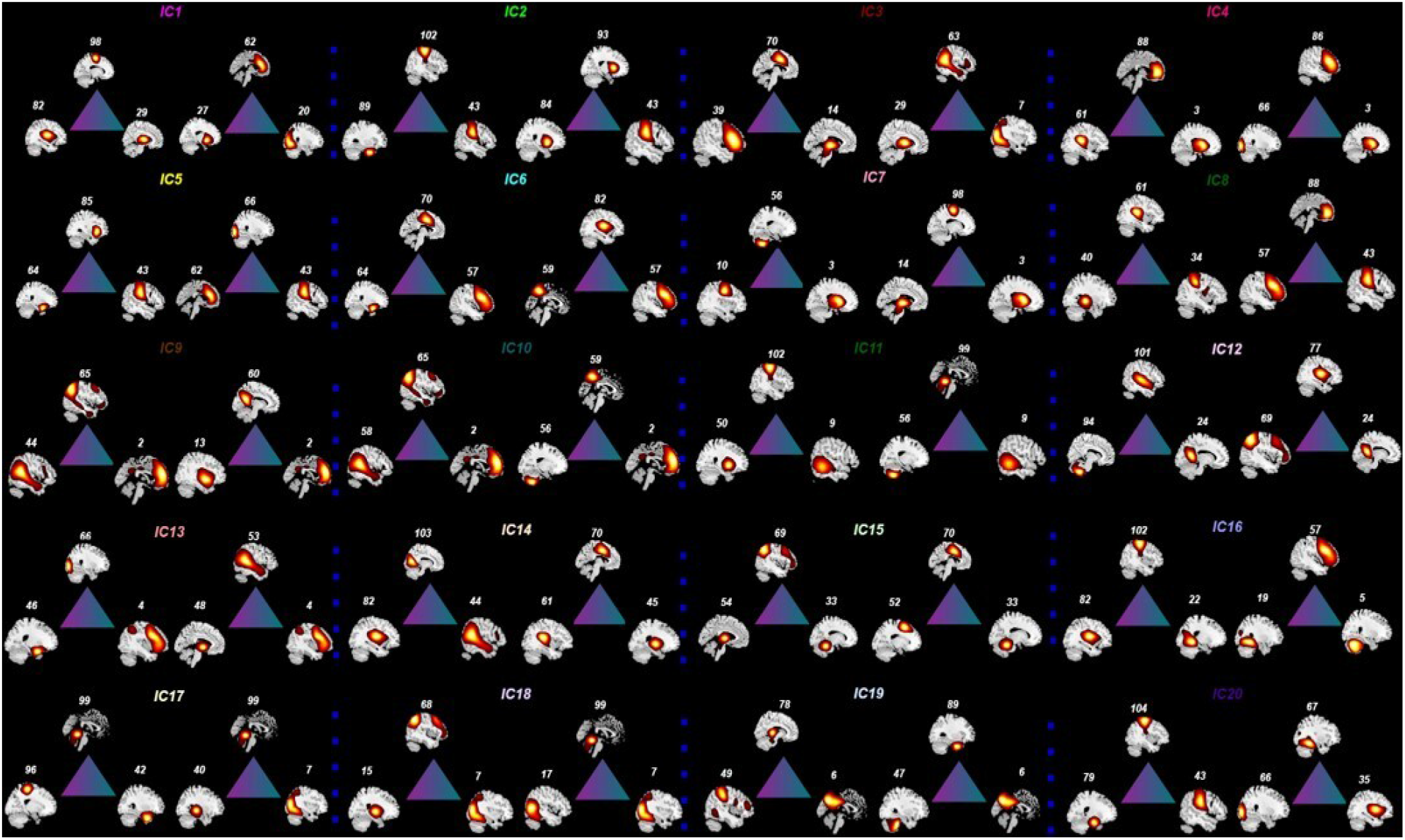
Triple spatial interaction components from GICA.

**Figure 3:**
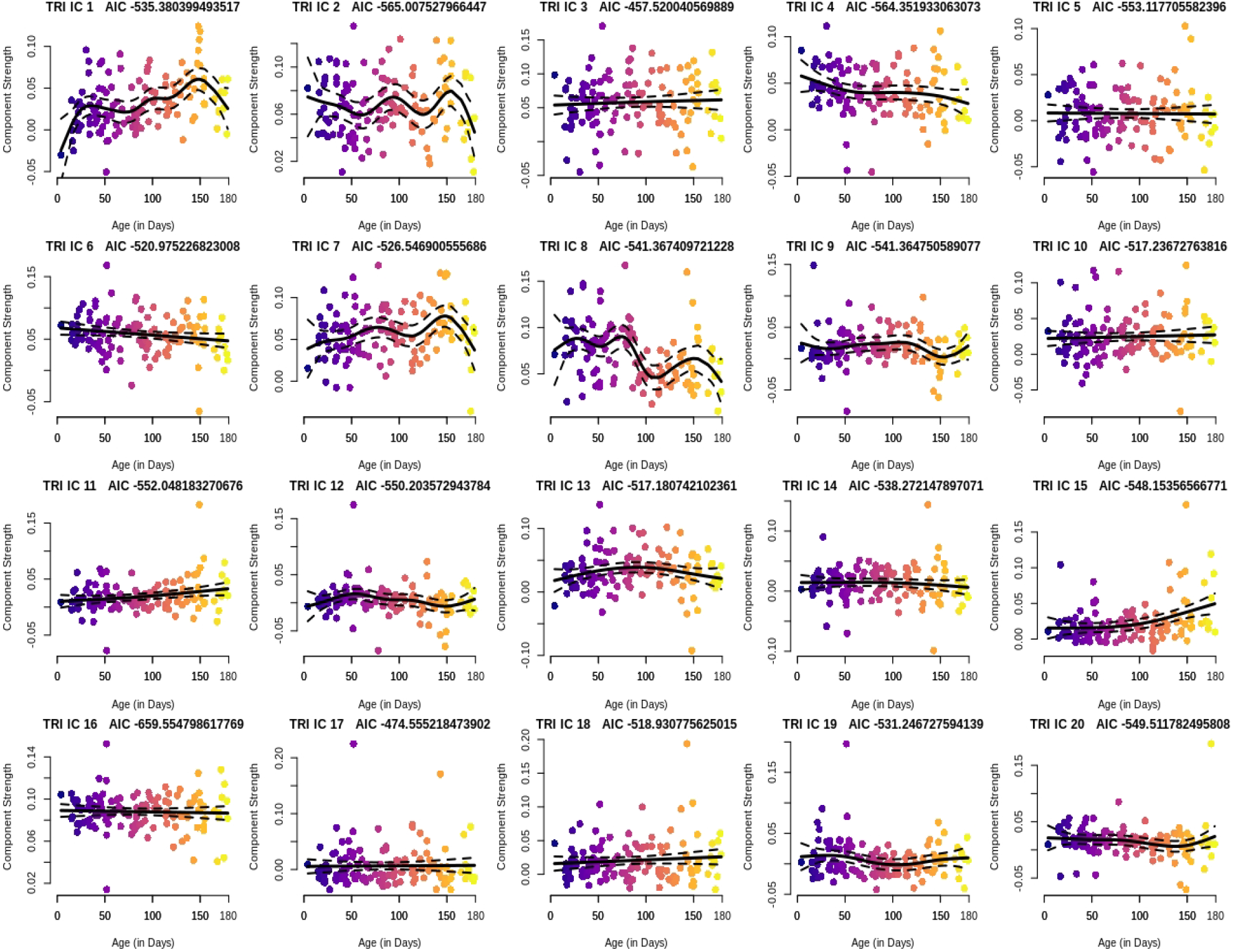
Age association with triple interaction components from GICA.

**Figure 4:**
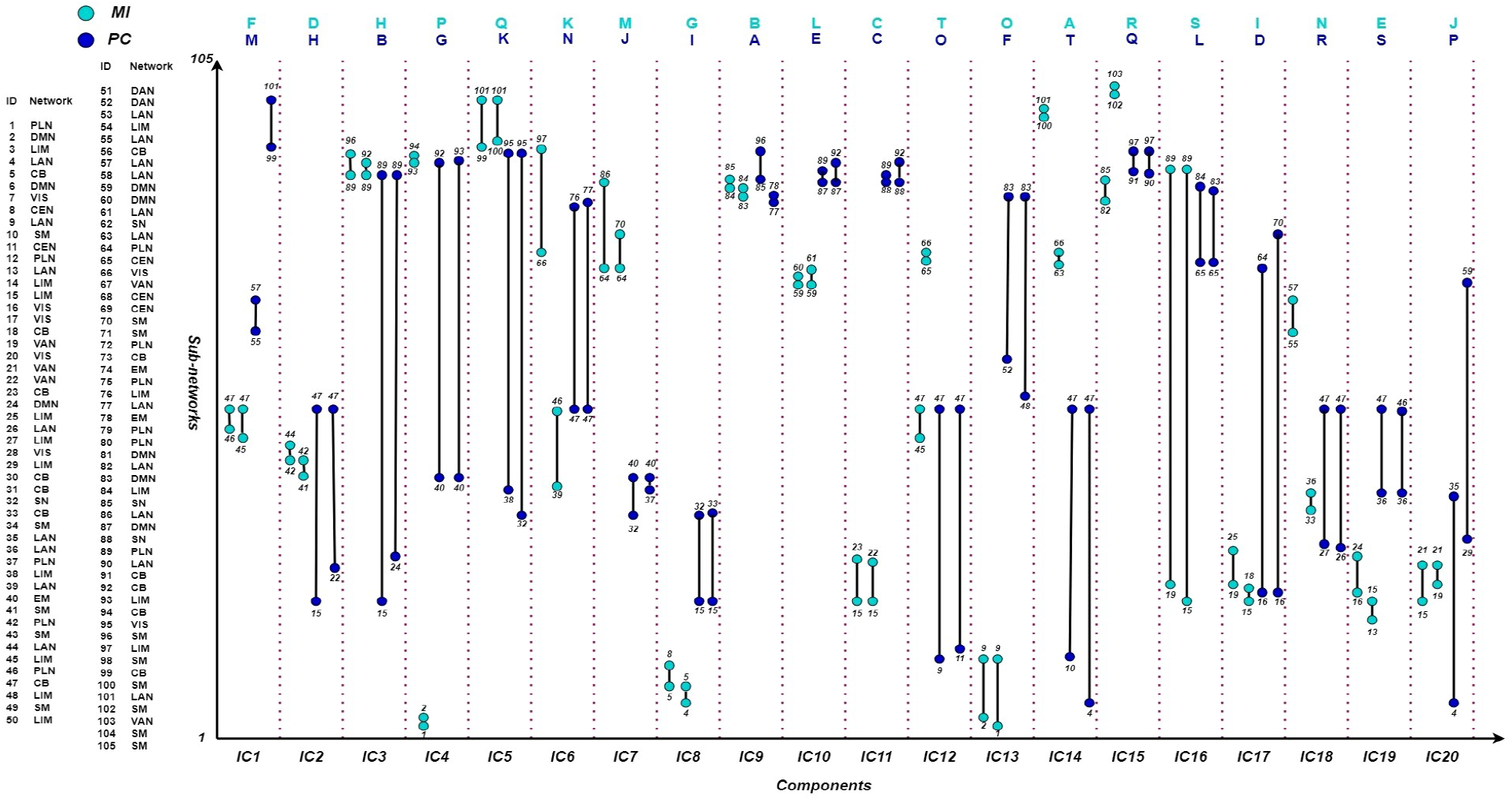
MI&PC interaction components from GICA. The order of the alphabet refers to network interaction strengths.

**Figure 5:**
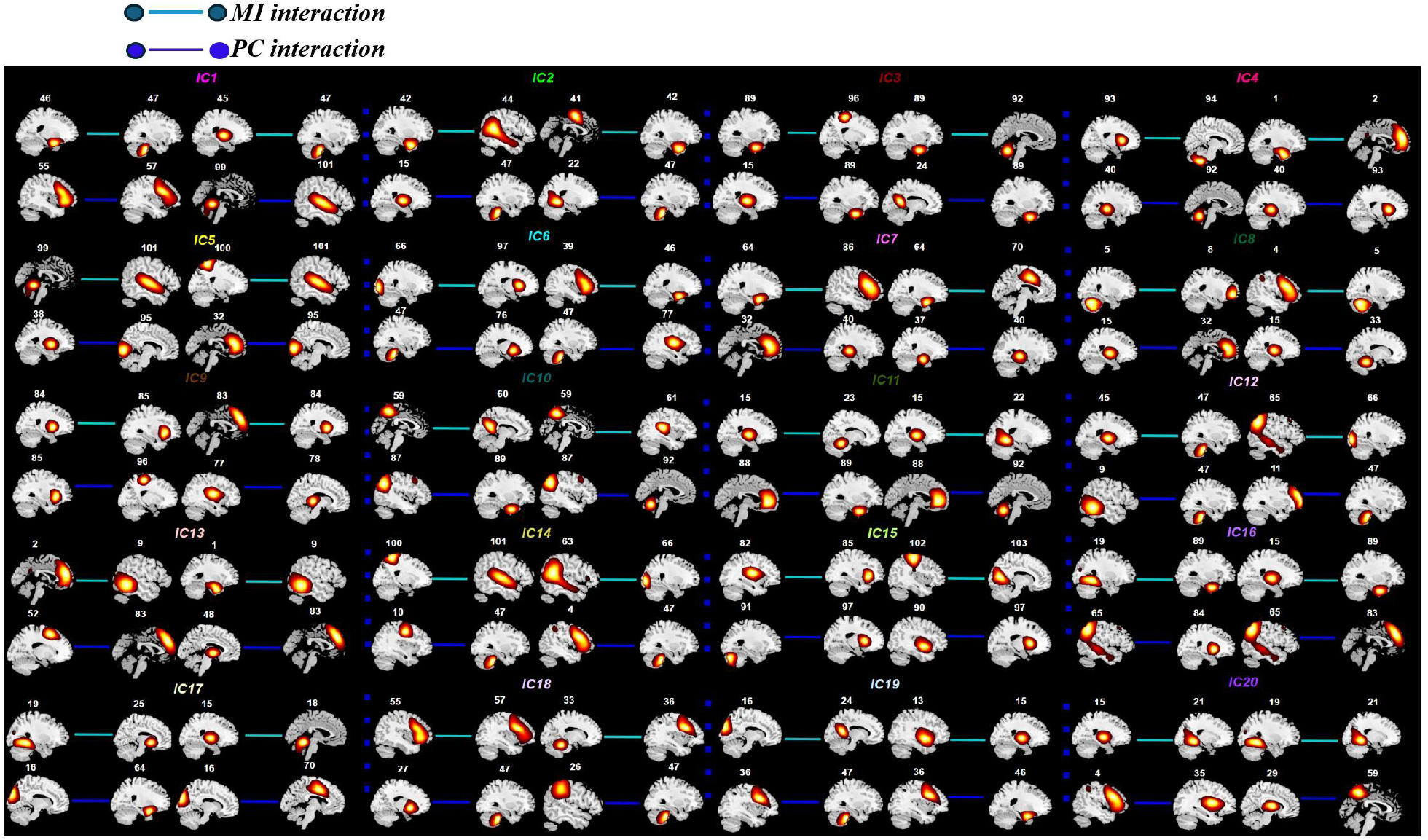
MI&PC spatial interaction components from GICA.

**Figure 6:**
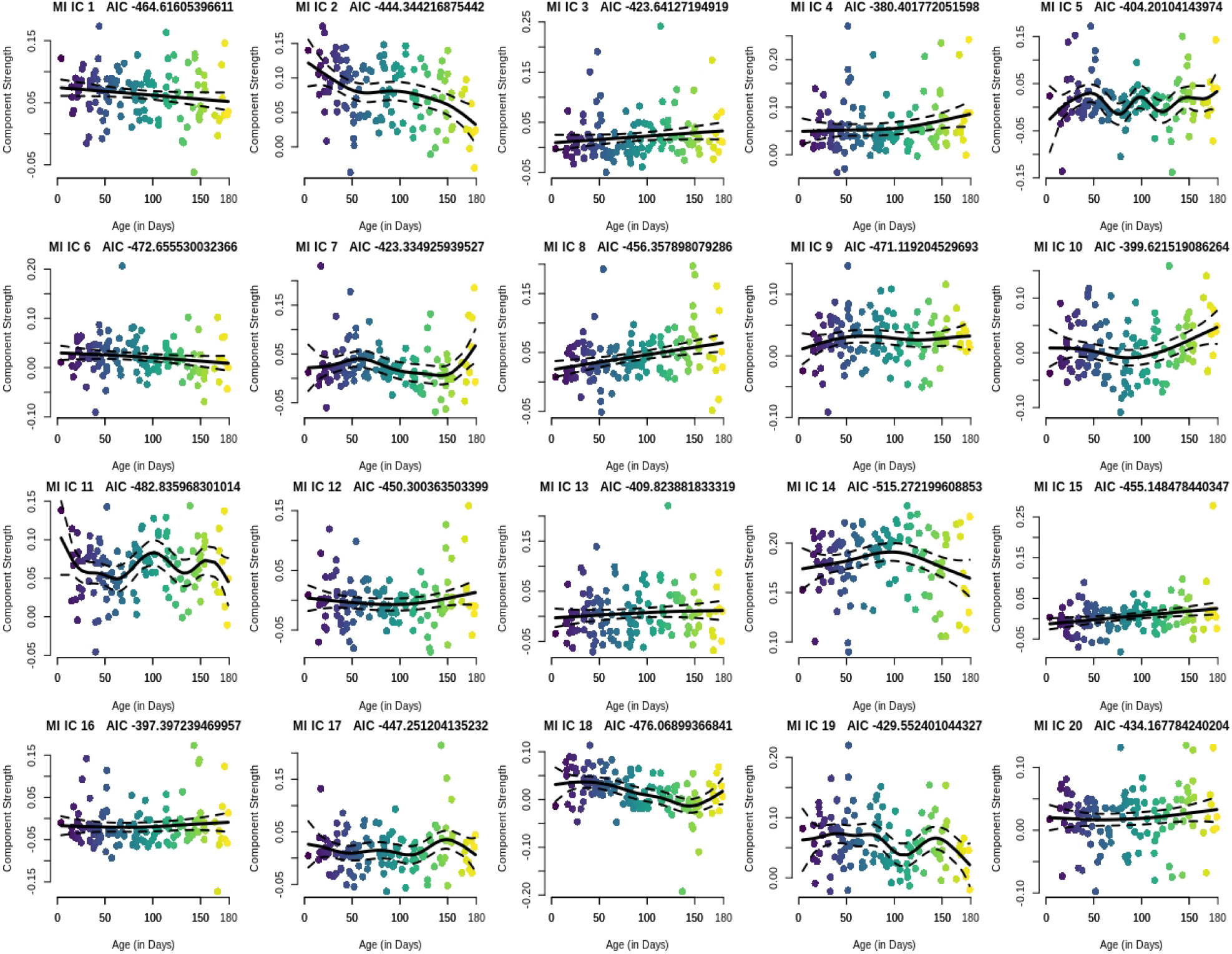
Age association with MI interaction components from GICA.

**Figure 7:**
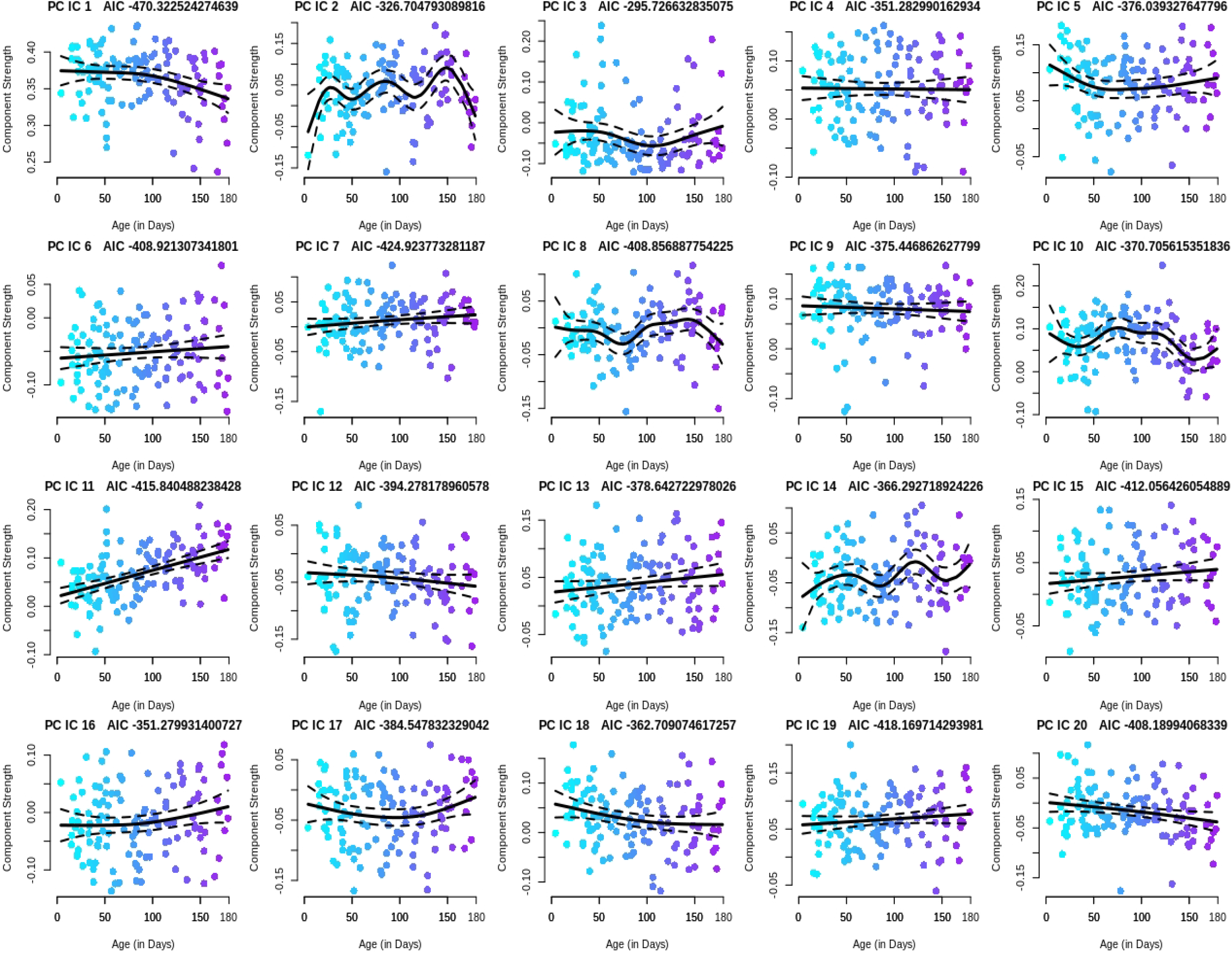
Age association with PC interaction components from GICA.

